# Estimating the effect of tissue- and blood-derived cell reference matrices on deconvolving bulk transcriptomic datasets

**DOI:** 10.1101/2025.01.24.634725

**Authors:** Siqi Sun, Shweta Yadav, Mulini Pingili, Dan Chang, Jing Wang

## Abstract

**Background:** Cell deconvolution is a method used to characterize the composition of the mixed cell population in bulk transcriptomic datasets. Tissue- and blood-derived cell reference matrices (CRMs) are typically used, but their impact on deconvolution has yet to be evaluated.

**Methods:** Recursive feature elimination and random forest methods were applied to build tissue- and blood-derived CRMs using single-cell RNA sequencing (scRNA-seq) datasets of inflammatory bowel disease (IBD) which comprises two subtypes (Crohn’s disease (CD) and ulcerative colitis (UC)). Combining with three published blood-derived CRMs (IRIS, LM22, and ImmunoStates), public bulk transcriptomes datasets and simulated datasets generated from scRNA-Seq were used to evaluate the deconvolution performance by goodness-of-fit and cell fractions correlation. Additionally, two infliximab-treated bulk datasets were used to compare the treatment-related cell types revealed by tissue- and blood-derived CRMs. Lung adenocarcinoma (LUAD) single-cell and TCGA bulk transcriptomic datasets were also used for evaluation.

**Results:** For CD and UC tissue bulk datasets, tissue-derived CRMs showed better deconvolution goodness-of-fit scores compared to blood-derived CRMs. Meanwhile, tissue-derived CRMs represented more accurate cellular proportion estimates for most cell types, such as immune and stromal cells in tissue pseudobulk datasets. They also revealed more treatment-related cell types. Additionally, CRMs derived from the scRNA-seq datasets from CD and UC colon samples had similar performance. In contrast, CRMs derived from CD ileum scRNA-seq dataset had better performance in the ileum bulk datasets. All CRMs yielded consistent deconvolution performance when deconvolving blood bulk transcriptomics. The similar results have also been shown using LUAD datasets.

**Conclusions:** Our results emphasize the importance of selecting appropriate CRMs for cell deconvolution, particularly in bulk tissue transcriptomes in immunology and oncology. Such considerations can be extended to encompass other disease implications.

**AbbVie Disclosure statement:** All authors are current employees of AbbVie. The design, study conduct, and financial support for this research were provided by AbbVie. AbbVie participated in the interpretation of data, review, and approval of the publication.

**Key Points:**

- Tissue-derived CRMs showed higher goodness-of-fit compared to blood-derived CRMs for deconvolving bulk tissue transcriptomics.
- All CRMs yield consistent goodness-of-fit for deconvolving bulk blood transcriptomics.
- Tissue-derived CRMs represent more accurate cellular proportion estimates and reveal more treatment-related cell types, and the specific tissue type is relevant to the deconvolution performance.

## Introduction

Bulk transcriptomic datasets have provided a comprehensive view of genome-wide transcriptomic variations in different conditions by measuring the average expressions across a large population of cells. However, bulk datasets cannot characterize the variation of cell type composition across subjects, which can provide insights into disease mechanisms and potential therapeutic targets. To address this challenge, cell deconvolution methods have been developed to estimate the relative abundance of individual cell types in each sample of the bulk transcriptomic dataset [1–3]. One type of widely used deconvolution methods is based on a predefined cell reference matrix (CRM) generated by human peripheral blood mononuclear cells (PBMCs) microarray datasets (e.g. IRIS [4], LM22 [5,6], and ImmunoStates [7]). CRMs were designed to capture genes that distinguish different hematopoietic cell types, with the potential to fit most deconvolution cases [7,8]. Based on this advancement, recent studies have identified cell types for various diseases and revealed potential treatment outcomes, such as cancer and autoimmune disorders [9–13]. For example, Thorsson et al. identified six immune subtypes spanning multiple tumor types and further characterized them with therapeutic and prognostic implications by applying deconvolution analysis of bulk RNA-seq data [12]. However, attention has primarily been focused on changes of the immunological cell types in the tissue microenvironment as CRMs were not well-established for other cell types that are not present in the peripheral blood [14]. It remains unclear to which extent the gene expression profile of tissue-derived cell types changes (e.g., epithelial and stromal cells) across various tissues or disease progression.

The rapid development of single-cell RNA sequencing (scRNA-seq) has advanced our understanding of the heterogeneity associated with immune and other cell types by profiling cell type-specific transcriptome, especially for the tissue samples. However, scRNA-seq still faces obstacles to retaining enough sample sizes and levels of profiles due to sample preservation requirement, labor-intensiveness and high cost, which prohibits its wide application in disentangling changes in cell frequencies [15,16]. Recent studies on CRMs derived from scRNA-seq have demonstrated an improved performance in leukocyte deconvolution [8] and expanded the possibilities for deconvolving diverse tissues, including the dorsolateral prefrontal cortex [17]. The crucial impact of appropriate and representative CRMs on deconvolution performance has been thus widely recognized [7,18,19]. However, there is a lack of comprehensive understanding of the effects of tissue- and blood-derived CRMs on deconvolving bulk transcriptomic datasets.

In this study, we first evaluated the effects of tissue- and blood-derived CRMs on cell deconvolution in Crohn’s disease (CD) and ulcerative colitis (UC). Using the public scRNA-seq datasets, we generated four CRMs (three tissue-derived CRM from colon and ileum and one blood-derived CRM from PBMCs). With three existing blood matrices, including IRIS, LM22, and ImmunoStates, we evaluated the deconvolution performance on public bulk transcriptomic and pseudobulk datasets. We further estimated the relationship between cell fractions and clinical information in the bulk transcriptomic with treatment information. To test whether the conclusion can be extended to other disease area, scRNA-seq and TCGA bulk transcriptomic datasets from lung adenocarcinoma (LUAD) were used to create lung-based CRM and compare with existing blood matrices.

## Methods

### Dataset collection and pre-processing

We searched Gene Expression Omnibus (GEO) [20] and ArrayExpress [21] for available CD or UC bulk transcriptomic studies until October 2023 and selected 44 datasets with at least ten patients (Fig. 1). In addition, we incorporated 600 lung biopsies from patients with lung adenocarcinoma (LUAD) in TCGA. Thus, our analyses included 1,375 blood, 1,779 colon, 1,154 ileum and 600 lung tumor samples (Table. S1). We have five different types of datasets: colon biopsies in CD patients (CD-colon), ileum biopsies in CD patients (CD-ileum), colon biopsies in UC patients (UC-colon), blood samples in CD patients (CD-blood), blood samples in UC patients (UC-blood) and lung biopsies in LUAD patients. Samples were sequenced in different microarray and RNASeq platforms (Table. S1). For IBD-related datasets, all data matrices had been downloaded from the respective original publications for microarray datasets and GEO for microarray and RNA-Seq datasets. In addition, LUAD RNA-seq datasets were downloaded from https://portal.gdc.cancer.gov/. Microarray data underwent RMA normalization, while RNA-seq data were normalized using the TMM method [23,24]. The gene-level expression for each sample was calculated by averaging expression values from probes mapping to the same genes while excluding individual probes that were associated with more than one transcript.

**Fig. 1.**
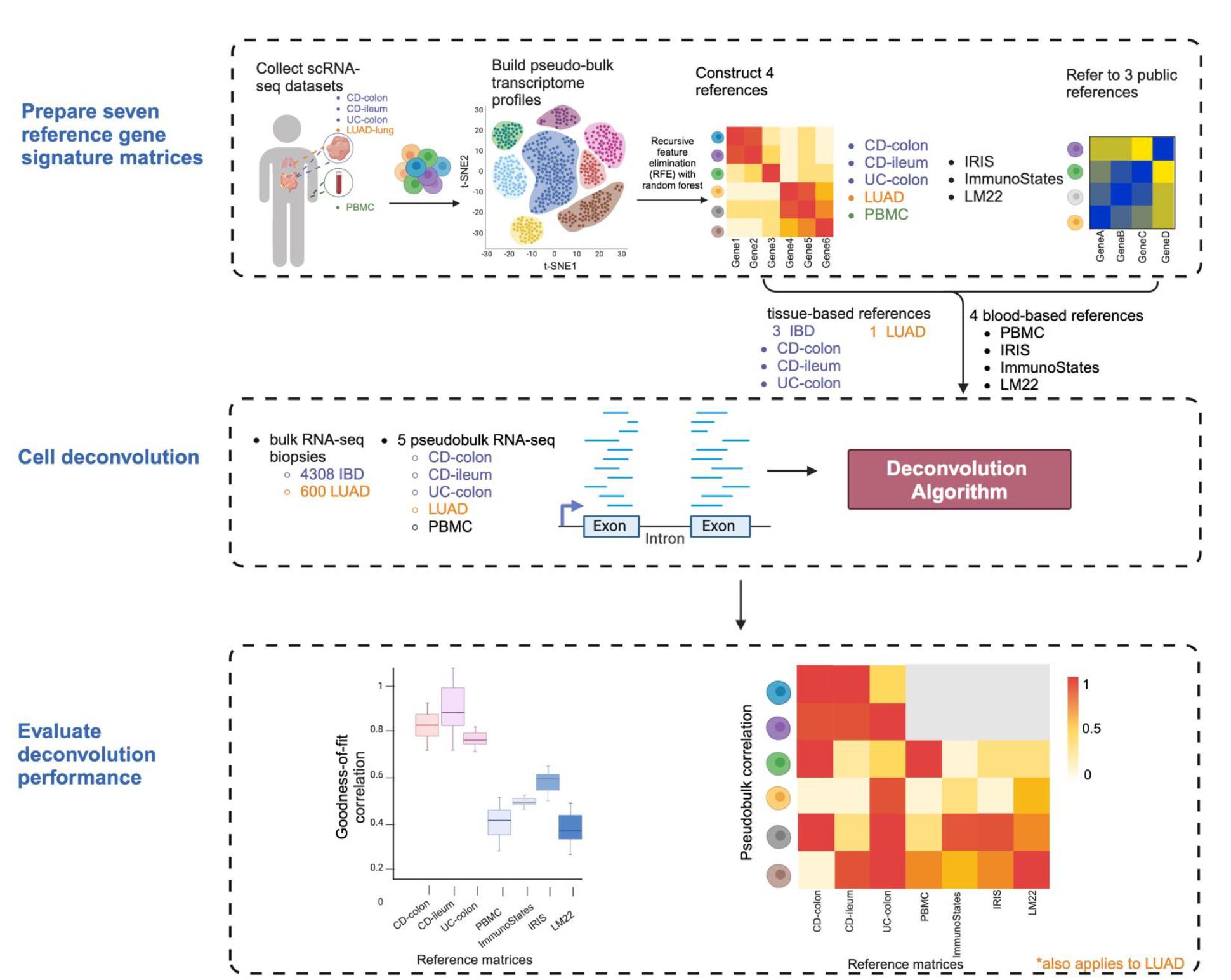
Analysis workflow. We first prepared eight reference matrices for our deconvolution analyses, including five generated from scRNA-seq and three public matrices. The deconvolution performance was evaluated by goodness-of-fit on 4908 bulk transcriptomic samples and cell fraction correlation on five pseudobulk datasets across immunology and oncology biopsies. IBD: inflammatory bowel disease; LUAD: lung adenocarcinoma; CD: Crohn’s disease; UC: ulcerative colitis; PBMC: human peripheral blood mononuclear cells; CD-colon: colon biopsies from CD patients; CD-ileum: ileum biopsies from CD patients; UC-colon: colon biopsies from UC patients; CD-blood: blood samples from CD patients; UC-blood: blood samples from UC patients. Created with BioRender.com.

The CD scRNA-seq datasets consist of colon and ileum biopsies with 45 and 58 cell types, respectively [25]. Colon biopsies from UC individuals with 51 cell types were also included [26]. The LUAD scRNA-seq dataset included lung biopsies with 10 cell types [27]. To identify whether the difference between tissue- and blood-derived CRM were from the platform difference (scRNA-seq vs microarray), we also included one PBMC scRNA-seq from CD individuals with 12 cell types [28]. Several QC on the scRNA-seq were further performed, including removing cells with abnormal mitochondrial RNA and gene levels (percent. mt < 25 & nFeature_RNA > 100 & nFeature_RNA < 5000), followed by keeping cell types with more than 500 cells (150 cells for LUAD). Further, genes were removed when the expression was limited to a small number of cells (less than 20% of cells in the smallest cell types of each scRNA-seq dataset), or the names related to HLA, immunoglobulin, RNA, MT, and RP based on HUGO gene names.

### CRMs creation and deconvolution

To create CRMs from scRNA-seq, pseudobulk datasets for five scRNA-seq datasets were first generated by summing counts by cell types to the sample level [29]. Then, differentially expressed genes (DEGs) of each cell type were identified by the Limma/Voom method [30]. The top 50 DEGs of each cell type were selected for the feature selection by the recursive feature elimination (RFE) method with a random forest model [24]. The optimal number of markers of each cell type was determined as the minimum with at least 0.9 AUROC (area under the receiver operating characteristic curve). The minimum number of markers for each cell type was set as 5, which provided at least 0.95 AUROC in most cell types for five datasets (Fig. S1). This method was iterated for each cell type for each pseudobulk dataset. Then, CRM was derived by calculating the mean expression of markers in each cell type in the five pseudobulk datasets. This resulted in four CRMs based on tissues (CD-colon, CD-ileum, UC-colon and LUAD) and one blood-derived CRM PBMC (Table. S2-S6). Three public blood-derived CRMs were also collected, including ImmunoStates, IRIS, and LM22. Specifically, the LM22 matrix was downloaded from the supplementary table [6], the IRIS matrix was from the CellMix R package, and the ImmunoStates matrix was from the IMMUNOSTATES website (https://khatrilab.stanford.edu/immunostates/) [31]. Then, SVR (support vector regression) was used to deconvolve the cell mixture [6].

### Evaluation of deconvolution performance across IBD bulk transcriptomes

Goodness-of-fit was applied to evaluate the deconvolution performance by different CRMs (Fig. 1). Briefly, for a given CRM **M** and a mixture sample gene expression vector **s**, deconvolution estimates the known cell proportion vector **p** such that

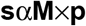

The goodness-of-fit of the matrix M is defined as the Pearson correlation coefficient between **s** and the reconstituted expression vector ŝ defined as **M** × **p** [6,7]. A higher goodness-of-fit score indicated good deconvolution accuracy.

### Simulated sample generation and evaluation

We generated 1,000 simulated bulk samples by randomly sampling cells without replacement from individual scRNA-seq datasets (Fig. 1). Each simulated sample contains 500 individual cells that were randomly selected from at least five cell types in one scRNA-seq subject (300 cells for LUAD from at least two cell types in LUAD) [18]. The cell type proportions were randomly chosen with a sum-to-one constraint. Afterward, the randomly selected single-cell expression profiles for each cell type were aggregated by summing their expression values. This random selection and aggregation process was iterated 1000 times to generate 1000 simulated samples [18].

To make comparisons between different CRMs, similar cell types were grouped into one cell lineage, considering that the initial generation and annotation of the single-cell transcriptome are from various sources (Table. 1). In this way, stromal cells were grouped into fibroblast and endothelial cells while epithelial cells were grouped into absorptive cells, secretory cells, and other epithelial cells. Additionally, immune cells were grouped into NK, mast cells, monocytes, B cells, CD4T, CD8T, and other lymphoid cells (Table. S7). The deconvolution performance was further evaluated by calculating the Pearson correlation coefficient between the estimated cell type proportion and the actual pseudobulk proportion across samples, specifically for each grouped cell lineage [7].

**Table 1.** Reference matrices.

| Reference matrix | Cell type number | Gene number | PMID references | Sources |
| --- | --- | --- | --- | --- |
| CD-colon | 45 | 217 | 31474370 | Constucted from single-cell datasets |
| CD-ileum | 58 | 282 | 31474370 |  |
| UC-colon | 42 | 199 | 31348891 |  |
| LUAD | 10 | 50 | 32822576 |  |
| PBMC | 12 | 69 | 31474370 |  |
| IRIS | 17 | 244 | 15789058 | Public |
| LM22 | 22 | 547 | 25822800 |  |
| ImmunoStates | 20 | 317 | 30413720 |  |

### Identification of treatment-related cell types

For IBD infliximab-treated datasets GSE16879 and GSE73661 (Table. S8), cell fractions were first estimated by different CRMs with SVR. Then, a p-value comparing cell fractions between responders and non-responders was calculated by Wilcoxon rank-sum and adjusted by the Benjamini-Hochberge method. Significant cell types were identified under the mean of cell fractions in at least one subtype>0.5%, and FDR<0.05.

### Data availability

GEO bulk datasets can be downloaded from public data portals. CD and UC scRNA-Seq data can be downloaded from single cell portal of the Broad institute (https://singlecell.broadinstitute.org/single_cell). The PBMC scRNA-seq dataset was downloaded from the website (https://scdissector.org/martin/) [28]. The processed LUAD scRNA-seq dataset was downloaded from Chan Zuckerberg Biohub Network (https://github.com/czbiohub-sf/scell_lung_adenocarcinoma)

All the analyses and plots were generated using the R programming language.

## Results

We first estimated the effect of CRMs on cell deconvolution in CD and UC by goodness-of-fit and the cell fraction estimates. Several treatment-related cell types were identified across different CRMs. Then the analyses were extended to LUAD for further evaluation and validation (Fig. 1).

### Establishment of CRMs based on single-cell transcriptomes in IBD

Using single-cell transcriptomes from CD and UC, we built CRMs for CD-colon, CD-ileum, UC-colon, and PBMC (Fig. 1). The mean AUROC of all cell types in each CRM was over 0.95 (Means±SDs: CD-colon 0.97±0.02, CD-ileum 0.98±0.02, UC-colon 0.98±0.02 and PBMC 0.98±0.03). This optimal performance can also be observed from the expression heatmap, in which selected maker genes had higher expression and were unique in each cell type (Fig. 2A-2D). Specifically, the CD-colon CRM comprises 217 genes from 45 different cell types, the CD-ileum CRM comprises 282 genes from 58 different cell types, the UC-colon matrix comprises 189 genes from 42 different cell types, and the PBMC CRM comprises 69 genes from 12 immune cell types (Table. 1). Notably, CD-colon and UC-colon CRMs share 75 common makers (p<2.2E-16) and CD-ileum and UC-colon CRMs share 72 common makers (p<2.2E-16) (Table. S2-S5, Fig. 2E), demonstrating that UC and CD may share similar etiology [32–34].

**Fig. 2.**
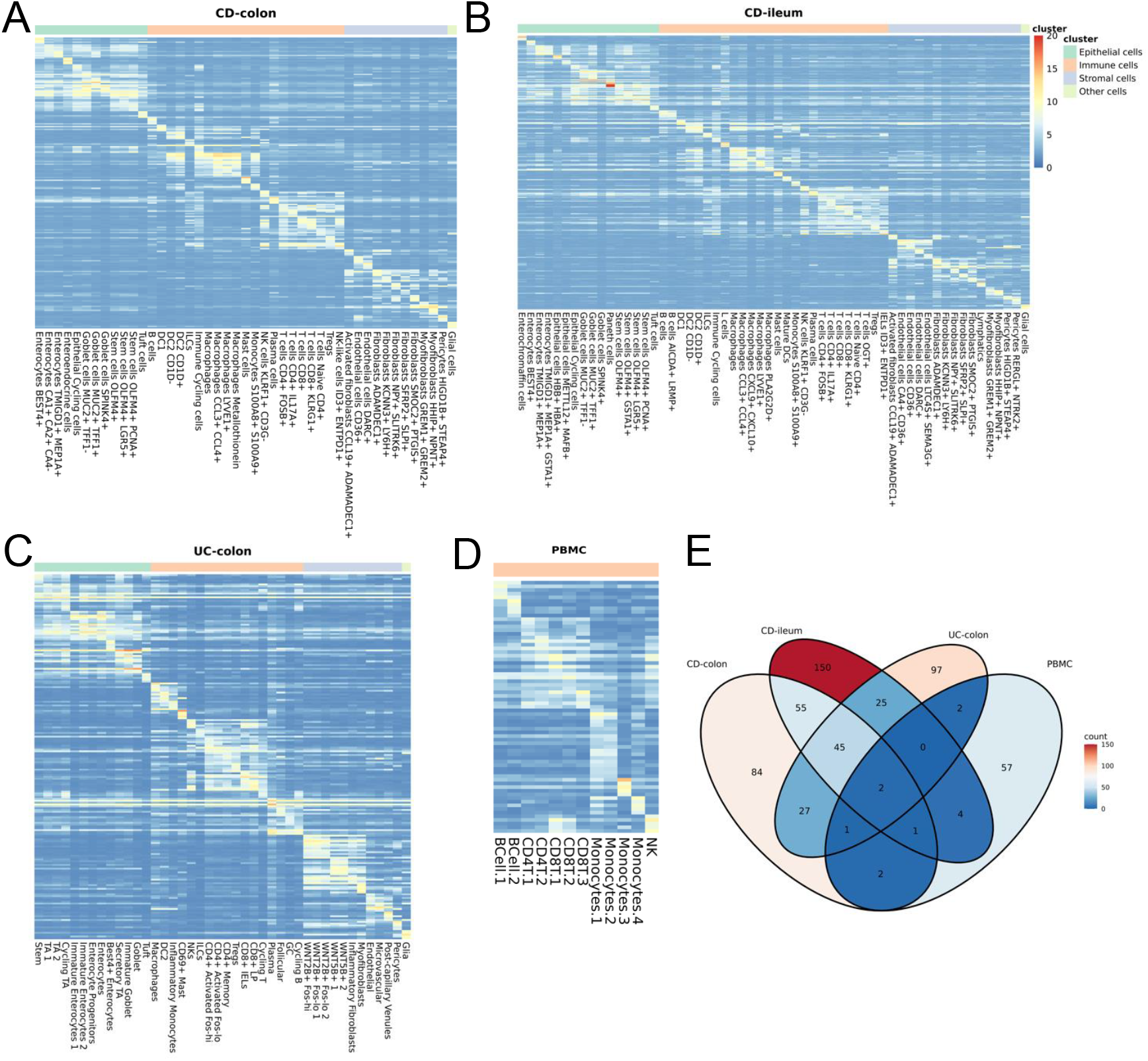
The selected makers were unique in each cell type for the generated reference matrices (A) CD-Colon (B) CD-ileum (C) UC-colon (D) PBMC. (E) Common markers between reference matrices. Tissue-based matrices exhibited a significant overlap in common markers with each other, while only a few markers were shared with the PBMC matrix.

### CRMs affect deconvolution performance

For the 44 independent public bulk transcriptomic datasets used in this study, we included a total of 796 samples from 10 CD-colon datasets, 1,154 samples from 7 CD-ileum datasets, 983 samples from 16 UC-colon datasets, 760 samples from 6 CD-blood datasets, and 615 samples from 5 UC-blood datasets (Fig. 1, Table. S1). For the ten CD-colon datasets, the goodness-of-fit for the three tissue-derived CRMs reached a minimum of 0.8. In contrast, the four blood-derived CRMs achieved less than 0.5 (Means±SDs: CD-colon 0.86±0.03, CD-ileum 0.79±0.03, UC-colon 0.85±0.04, PBMC 0.35±0.07, ImmunoStates 0.43±0.06, IRIS 0.46±0.04, LM22 0.35±0.09) (Fig. 2, Table. S9). The drastic differences were also observed in CD-ileum and UC-colon datasets (Fig. S2, S3, Table. S9), demonstrating higher accuracy for tissue-derived CRMs for deconvolving tissue datasets. On the other hand, all CRMs achieved a similar goodness-of-fit across bulk blood datasets (CD-blood and UC-blood) (Means±SDs: CD-colon 0.79±0.04, CD-ileum 0.84±0.03, UC-colon 0.76±0.05, PBMC 0.75±0.07, ImmunoStates 0.64±0.05, IRIS 0.57±0.05, LM22 0.56±0.04) (Fig. 4, Table. S9).

Typically, the deconvolution performance would be optimal with reduced bias when the matrix is specifically designed for each tissue [8]. However, we observed the trend of similar deconvolution performance using three tissue-derived matrices on deconvolving all 33 tissue bulk datasets (Means±SDs: CD-colon 0.83±0.04, CD-ileum 0.80±0.04 and UC-colon 0.80±0.06) (Fig. 2, S2, S3, Table. S9). Specifically, when deconvolving the CD-colon datasets, the goodness-of-fit of the CD-colon CRM is 0.86 compared to 0.79 achieved in the CD-ileum CRM (Table. S9). For the CD-ileum datasets, the deconvolution correlation of the CD-colon CRM is 0.81 compared to 0.83 achieved in CD-ileum (Table. S9). This suggests the deconvolution correlation of a tissue-derived CRM did not favor the subtypes of diseases (UC or CD) or affected sites (colon or ileum) (Fig. 2, S2, S3, Table. S9) [27–29].

More importantly, the deconvolution performance was unbiased toward the platforms and technologies of the bulk datasets being deconvolved and the matrix used for building. For example, when deconvolving the blood-derived datasets from microarray, the blood-derived CRMs built from microarray (ImmunoStates, IRIS, and LM22) yielded a goodness-of-fit similar to that of in PBMC from scRNA-Seq (Means±SDs: PBMC 0.75±0.1, ImmunoStates 0.70±0.04, IRIS 0.59±0.07, LM22 0.61±0.03). Similarly, the minimal differences in goodness-of-fit between different CRM platforms can also be observed in blood-derived datasets from RNA-seq (Means±SDs: PBMC 0.74±0.05, ImmunoStates 0.61±0.01, IRIS 0.57±0.02, LM22 0.54±0.01) (Table. S9). Taken together, the scRNA-Seq-derived CRM were not outperformed by the microarray-derived CRM for cell deconvolution (Fig. 3).

**Fig. 3.**
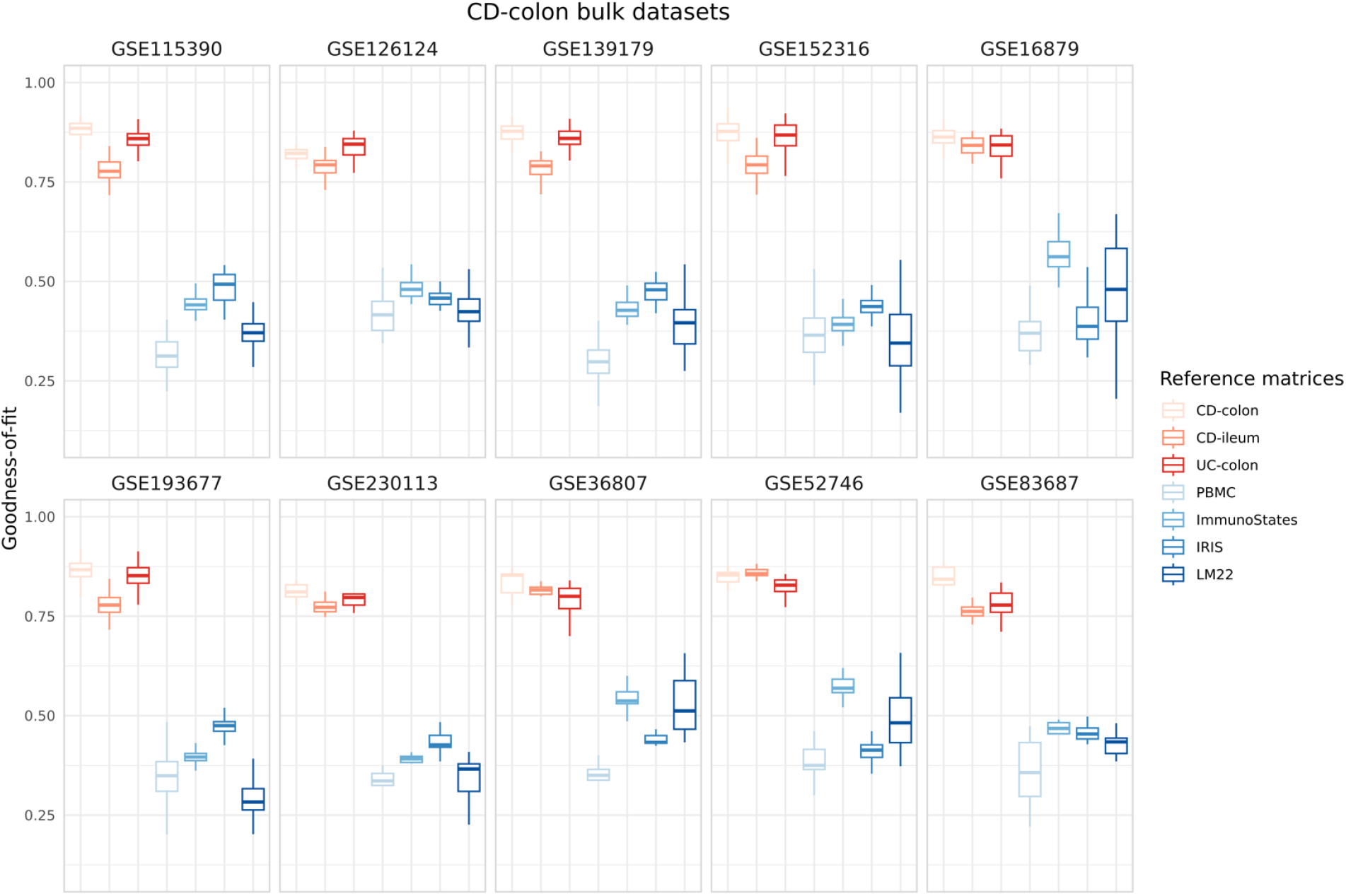
Deconvolution performance on 10 public CD-colon bulk transcriptomic datasets. The goodness-of-fit of each sample in the dataset was measured with different matrices. The results were shown using a whisker plot. Tissue-based matrices were depicted in red, while blood-based matrices were represented in blue. The individual dataset information can be found in the Table. S5.

**Fig. 4.**
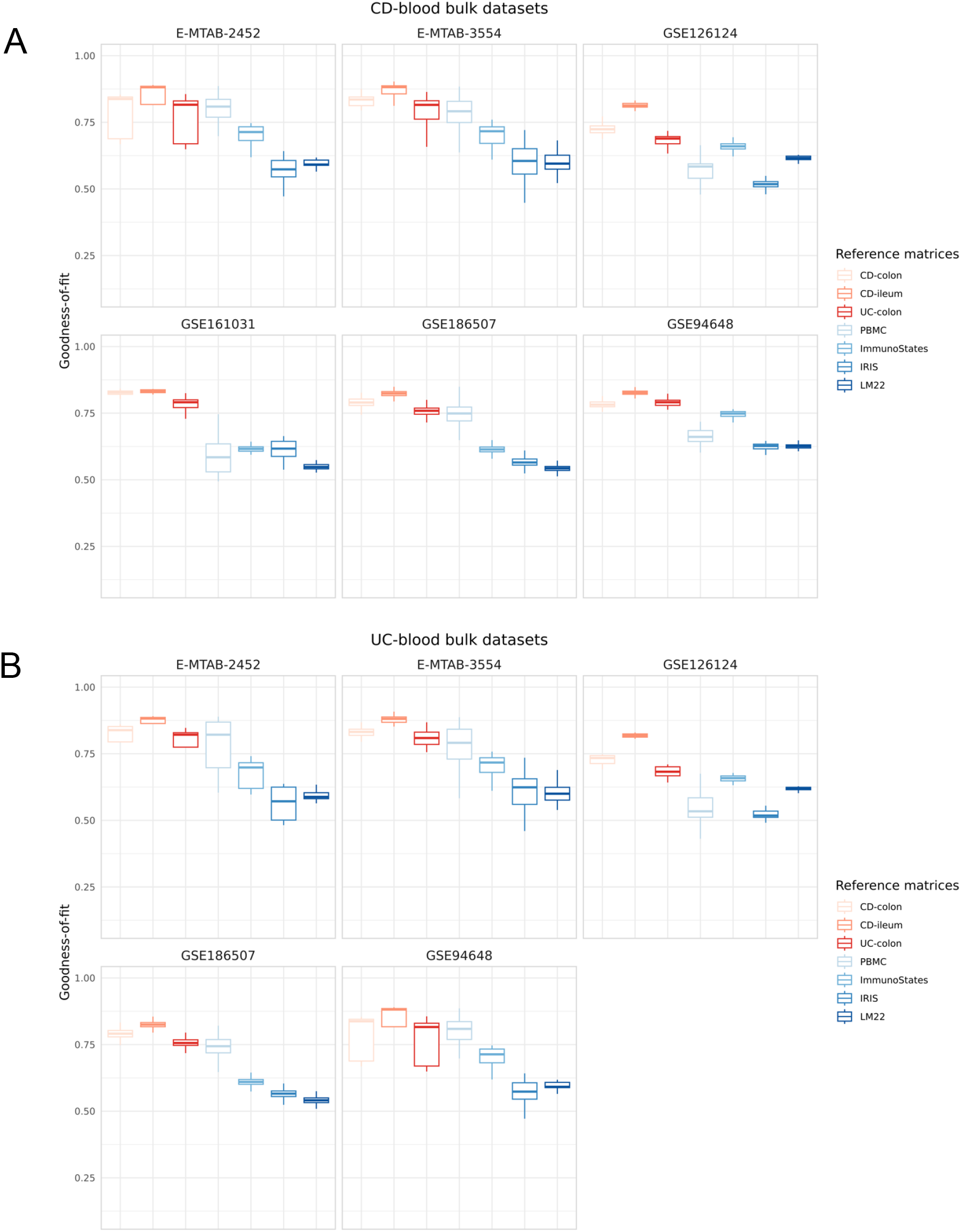
Deconvolution performance on public (A) CD-blood (B) UC-blood bulk transcriptomic datasets.

### The tissue-derived CRMs reflect representative cell fraction estimates in simulated samples

To further evaluate the performance, we built simulated samples for each single-cell dataset of colon, ileum and PBMC, and estimated the correlation of cell lineage proportions from ground truths and cell deconvolution (Fig. 1). As expected, the CRMs performed the best when deconvolving the matched simulated dataset (Fig. 5). As shown in the CD-colon simulated dataset, six cell lineages had high correlation (>0.7) in the CD-colon CRM compared to four in CD-ileum, and three in UC-colon (Fig. 5A). Correlations of at least 0.6 were achieved for the matched deconvolution (Means±SDs: CD-colon 0.62±0.20, CD-ileum 0.62±0.12 and UC-colon 0.67±0.17, PBMC 0.70±0.09). The tissue-derived CRMs showed a high correlation on epithelial and stromal cells, while the blood-derived CRMs failed to capture the signals (Fig. 5A). Across the three tissue-derived simulated datasets, for example, CD-colon achieved 0.8 for endothelial cells and 0.89 for glia; CD-ileum achieved 0.74 for endothelial cells and 0.85 for glia; UC-colon achieved 0.84 for endothelial cells and 0.87 for glia. For the immune cell estimates, tissue-derived CRMs also achieved higher correlation (Means±SDs: CD-colon 0.56±0.20, CD-ileum 0.60±0.16, UC-colon 0.60±0.18, PBMC 0.33±0.23, ImmunoStates 0.44±0.19, IRIS 0.32±0.19, LM22 0.47±0.23). Additionally, the estimation accuracy for the immune cell fractions was similar across CRMs in the PBMC simulated samples (Means±SDs: CD-colon 0.52±0.21, CD-ileum 0.57±0.19, UC-colon 0.57±0.15, PBMC 0.70±0.10, ImmunoStates 0.64±0.17, IRIS 0.53±0.14, LM22 0.65±0.18) (Fig. 5B), suggesting that all CRMs have captured representative immune markers for deconvolution (Table. S2-S5). Notably, the fractions of several immune cell types were consistently higher regardless of CRMs and simulated samples, including B cells and monocytes (Means±SDs: B cells 0.64±0.14, Monocytes 0.52±0.21) (Fig. 5B). The results evaluated by cell lineage estimates are consistent with goodness-of-fit and together provide strong evidence that the reference matrix is the major determinant of deconvolution accuracy, particularly for tissue-derived datasets. The overall performance is minimally affected by other factors such as platform, disease subtypes, and affected locations (Fig. 3, 4, S2, S3).

**Fig. 5.**
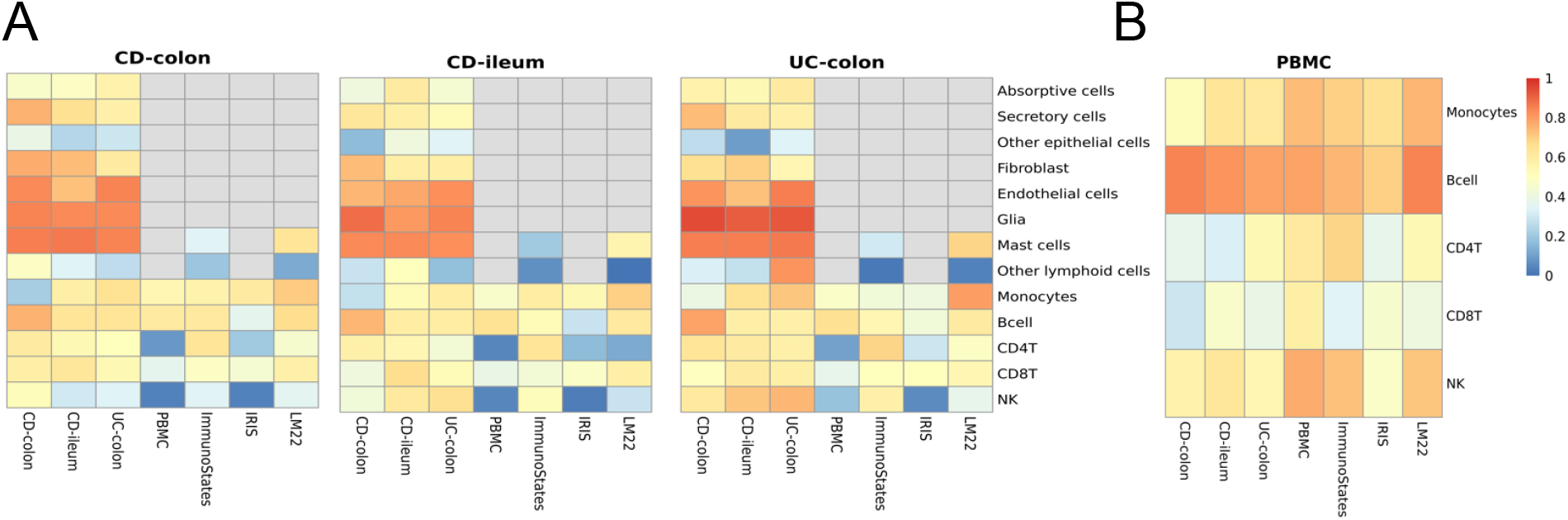
The correlation of cell lineage proportions on simulated (A) tissue and (B) blood pseudobulk datasets. The correlation between the estimated and ground truth in cell lineages was visualized using different colors to represent varying levels of correlation. High correlation was denoted by red, while low correlation was represented by blue. Grey was used to indicate missing values, as blood-based matrices were unable to capture some cell lineage.

### Tissue-derived CRMs revealed more treatment-related cell types

To identify cell types related to disease mechanisms and treatment response, we applied different CRMs to two infliximab-treated datasets and calculated significantly different cell types between responders and non-responders. For the GSE16879 dataset, the tissue-derived CRMs had the ability to discover the significant epithelial and stromal cells while maintaining the same sensitivity for identifying significant immune cells as blood-derived CRMs (FDR ≤ 0.05) (Fig. 6). For example, NK, CD4T and absorptive cells were identified using the CD-colon matrix whereas only immune cells, such as NK and mast cells were identified using ImmunoStates in the CD-colon dataset (Fig. 6A). The trend can also be observed in the GSE73661 UC-colon dataset with/without patient responses to infliximab or vedolizumab (Fig. 6D).

**Fig. 6.**
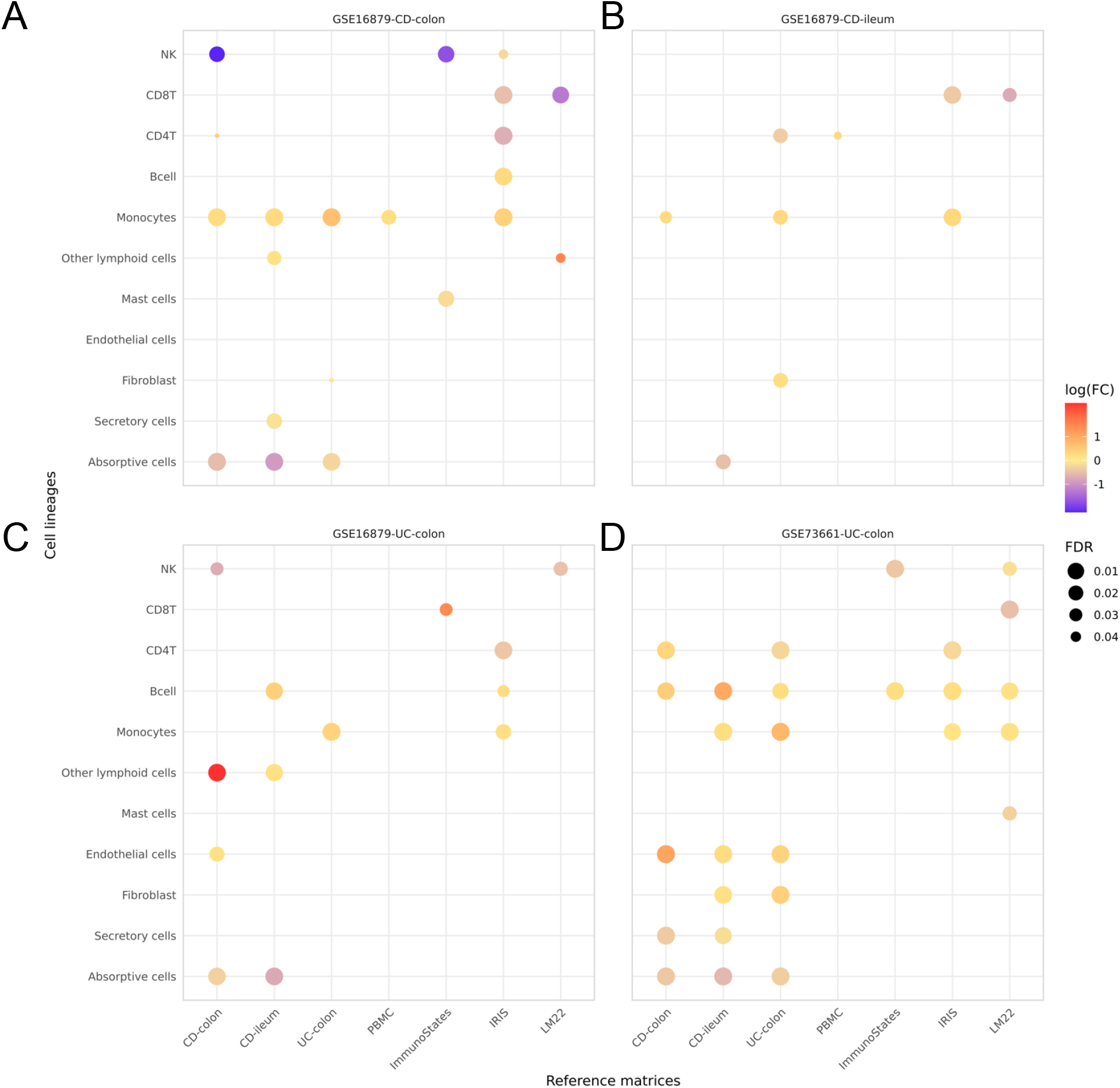
Tissue-based reference matrices identify more significant cell types between responders and non-responders in GSE16879 (A-C) and GSE73661 (D). The color of the log(FC) denotes the differences in cell fractions between non-responders and responders. Log(FC): log10 fold change; FDR: false discovery rate.

### LUAD CRM reveals more biological insights in lung

The LUAD CRM comprises 50 genes that differentiate 10 different cell types with high expression (Fig. 7A). Across 600 sample, the goodness-of-fit for the tissue-derived CRM (LUAD) reached at least 0.7 in two LUAD datasets. In contrast, the three blood-derived CRMs achieved less than 0.5 (Means±SDs: LUAD 0.78±0.06, ImmunoStates 0.41±0.03, IRIS 0.45±0.03, LM22 0.37±0.05) (Fig. 7B, 7C). When deconvolving LUAD simulated samples, LUAD CRM showed a high correlation on fibroblast, endothelial and epithelial cells, while the blood-derived CRMs failed to capture the signals (Fig. 7D). For the immune cell estimates, LUAD CRM also achieved higher correlation compared to the blood-derived CRMs (Means±SDs: LUAD 0.90±0.04, ImmunoStates 0.80±0.11, IRIS 0.67±0.14, LM22 0.75±0.11) (Fig. 7D). This result was consistent with the ones from CD and UC, indicated a general lessons of CRM selection across different diseases.

**Fig. 7.**
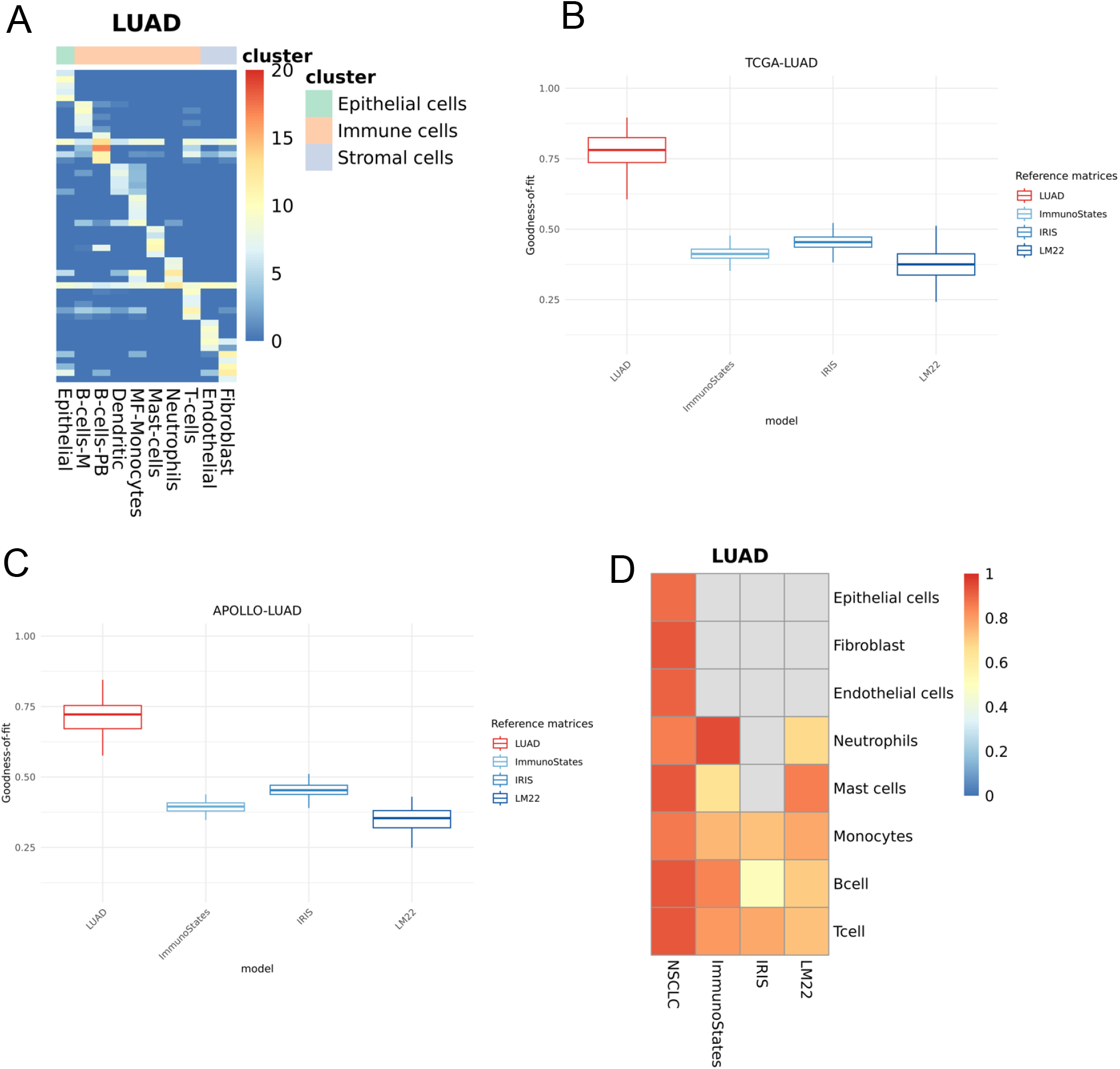
Deconvolution evaluation in LUAD. (A) LUAD reference matrix generated using sc-RNA seq (B, C) deconvolution performance evaluated based on goodness-of-fit with publicly available bulk transcriptomics datasets of LUAD lung samples (D) The correlation of cell lineage proportions on simulated LUAD pseudobulk datasets.

## Discussion

Knowledge of cell type compositions in disease-relevant tissues is an important step towards the identification of cellular targets of disease. Several computational methods have been developed to characterize the variation of cell type compositions [1–3]. These methods often rely on a CRM based on the pre-selected cell type-specific marker genes from tissue or blood datasets. Although many studies used blood-based (public) CRMs to deconvolve the tissue transcriptomic datasets[4–7,14], the effects of tissue- or blood-derived CRMs on cell deconvolution performance are not fully evaluated. Here, we performed comprehensive evaluations on deconvolution using goodness-of-fit and cell fraction estimates in public and simulated bulk transcriptomics datasets across autoimmune and oncological diseases. Our results showed that the source of the CRMs, whether derived from tissue or blood, needs to be considered for deconvolution, especially for bulk tissue transcriptomes. In contrast, blood bulk datasets are not sensitive to the specific CRMs. Other factors, including the sequencing platform used for CRM building (microarray or RNA-seq), disease subtypes (UC or CD), and affected locations (colon or ileum), have minimal effects on the overall performance. Additionally, the tissue-derived matrices reflected representative cell fraction estimates in simulated samples and revealed more treatment-related cell types between responders and non-responders.

One of the reasons that tissue-derived CRMs outperformed blood-derived CRMs in tissue datasets may be that genes used to estimate the cell type proportions are expressed in the cell types that are absent in public CRMs (e.g. epithelial and fibroblast cells). To investigate the deconvolution performance of immune-related cell types across different CRMs, we constructed tissue-derived CRMs that limited to immune-related cell types (CD-colon_imm, UC-colon_imm and CD-ileum_imm). This ensures that all CRMs have comparable immune cell types. As shown in Fig. S4 and S5, we observed that tissue-derived CRMs still showed higher goodness-of-fit compared to blood-derived CRMs for deconvolving bulk tissue transcriptomics. For example, when deconvolving CD-colon datasets, the goodness-of-fit of three tissue-derived immune-only CRMs reached a minimum of 0.7 (Fig. S4A, Table. S10, Means±SDs: CD-colon_imm 0.80±0.04, CD-ileum_imm 0.78±0.03, UC-colon_imm 0.74±0.03, PBMC 0.35±0.07, ImmunoStates 0.43±0.06, IRIS 0.46±0.04, LM22 0.35±0.09). The drastic differences were also observed in CD-ileum and UC-colon datasets (Fig. S4B, S5, Table. S10). This suggests that the immune cell proportion estimated from tissue-derived CRMs are more accurate than those from blood-based CRMs. This aligns with our findings from simulated samples with ground-truth composition from scRNA-seq datasets. These results proved our hypotheses and suggested that tissue bulk datasets should be deconvolved by the tissue-derived CRMs to gain comprehensive biological insights on both immune and non-immune cells in the tissue.

Notably, SVR was used as the benchmarking approach to evaluate deconvolution performance using different CRMs in this study. It has been shown that robust regression approaches, such as RLM, have given comparable results to SVR across different datasets [18]. We also observed similar performance for deconvolving IBD datasets using RLM (Robust Fitting of Linear Models) [35]. For example, the goodness-of-fit for the three tissue-derived CRMs reached a minimum of 0.75 when deconvolving CD-colon datasets. In contrast, the four blood-derived CRMs achieved less than 0.5 (Means±SDs: CD-colon 0.86±0.03, CD-ileum 0.78±0.04, UC-colon 0.85±0.03, PBMC 0.38±0.07, ImmunoStates 0.42±0.05, IRIS 0.47±0.04, LM22 0.35±0.08) (Fig. S6A, Table. S11). The drastic differences were also observed in CD-ileum and UC-colon datasets (Fig. S6B, S7, Table. S11), demonstrating higher accuracy for tissue-derived CRMs for deconvolving tissue datasets. On the other hand, all CRMs achieved a similar goodness-of-fit across bulk blood datasets (CD-blood and UC-blood) (Means±SDs: CD-colon 0.77±0.002, CD-ileum 0.82±0.01, UC-colon 0.76±0.001, PBMC 0.75±0.0002, ImmunoStates 0.61±0.001, IRIS 0.57±0.0004, LM22 0.56±0.001) (Fig. S8, Table. S11). Other newly developed algorithms were designed to account for cross-subject differences and batch-effect confounding, such as MuSiC (MUlti-Subject SIngle Cell deconvolution) [36]. MuSiC applies weighted non-negative least squares regression (W-NNLS) and does not require pre-determined CRMs and further applied for deconvolution for comparison [15]. With cell-type specific gene expression from tissue-derived scRNA-seq (CD-colon, CD-ileum and UC-colon), MuSiC showed a high correlation on epithelial and stromal cells (Means±SDs: CD-colon 0.65±0.26, CD-ileum 0.57±0.27, UC-colon 0.60±0.1) (Fig. S9), which was consistent with SVR and RLM results. However, MuSiC failed to deconvolve some immune cell types (Fig. S9) [37].

It has been suggested that the compromised function of tissues against microorganisms is a pivotal factor in the onset of IBD. This impairment can result from intrinsic dysfunction in epithelial, immune, and stromal cells or the compensatory actions of other cells attempting to restore homeostasis [38–40]. Similarly, in our analyses of tissue bulk datasets between non-responders and responders, we observed significant differences in immune cell types, such as monocytes, across all matrices (FDR ≤ 0.05) (Fig. 6). Furthermore, fibroblasts were found to be significantly different using the tissue-derived CRMs for deconvolution (Fig. 6B, 6D). This is consistent with previous studies regarding the important role of fibroblasts in achieving long-lasting remission with anti-TNF therapy in IBD patients [26,28]. Collectively, this suggests that analysis using tissue-derived CRMs detected significant epithelial and stromal cells for treatment responses, while matching results using blood-derived CRMs in immune cell sensitivity.

In this study, we evaluated the effects of CRMs on deconvolution accuracy using IBD and LUAD datasets. These results underscored important considerations for accurate deconvolution, such as the sources of the reference matrix and the tissue types of bulk transcriptomics. As bulk tissue data are more easily accessible than single-cell RNA-seq, an appropriate selection for reference matrix allows the utilization of the vast amounts of sequencing data for deepening cellular understanding of disease progression and treatment courses. Such insights may extend beyond IBD and LUAD to other diseases in immunology and oncology, offering valuable guidance applicable to various disease types across therapeutic areas.

## Author Biographies

All authors are current employees of AbbVie.

