## Supplementary material for "Estimating the effect of tissue- and blood-derived cell reference matrices on deconvolving bulk transcriptomic datasets": Fig. S1-Fig. S9

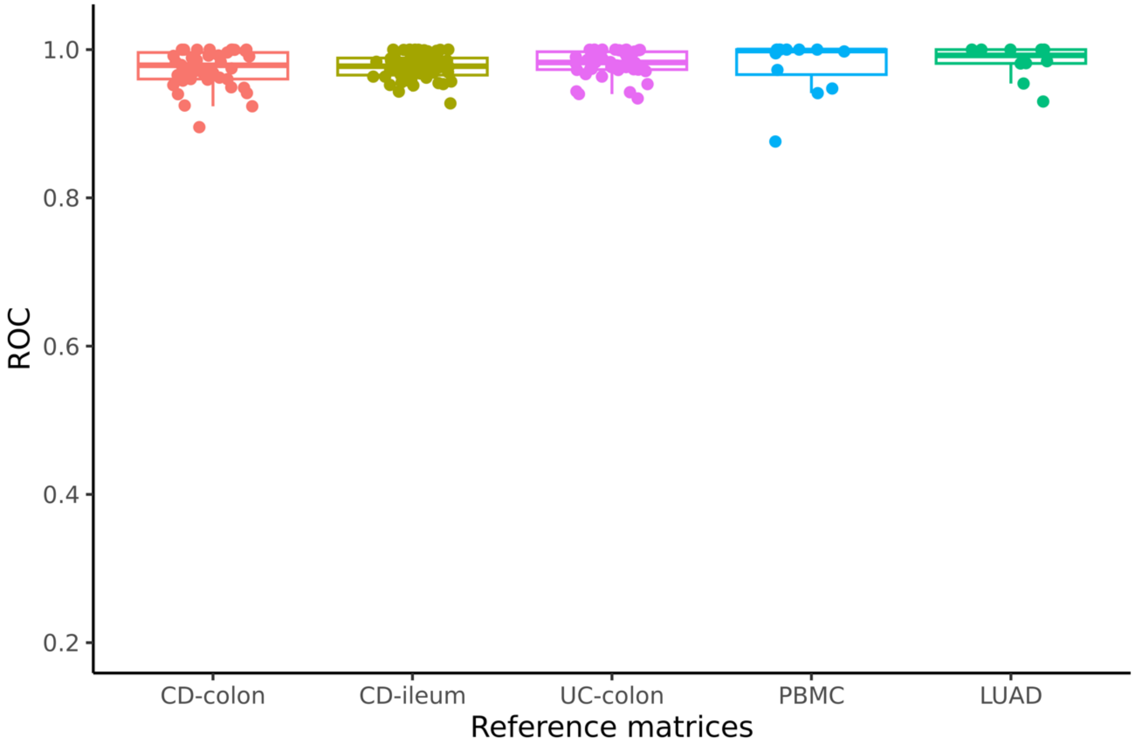


Fig. S1 The top 5 DEGs in most cell types achieved an AUROC score of greater than 0.95.

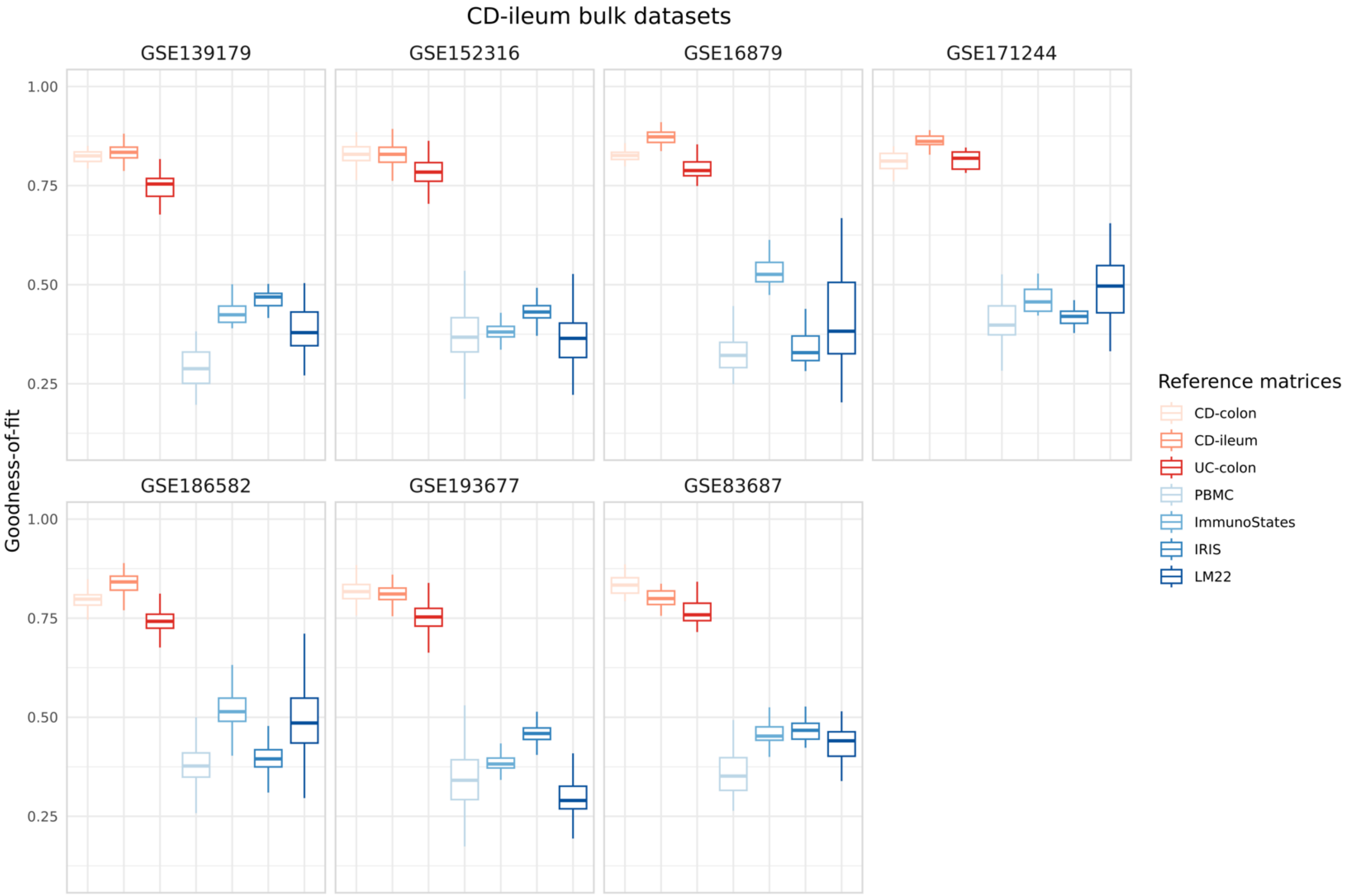


Fig. S2 Deconvolution performance on seven public CD-ileum bulk transcriptomic datasets.

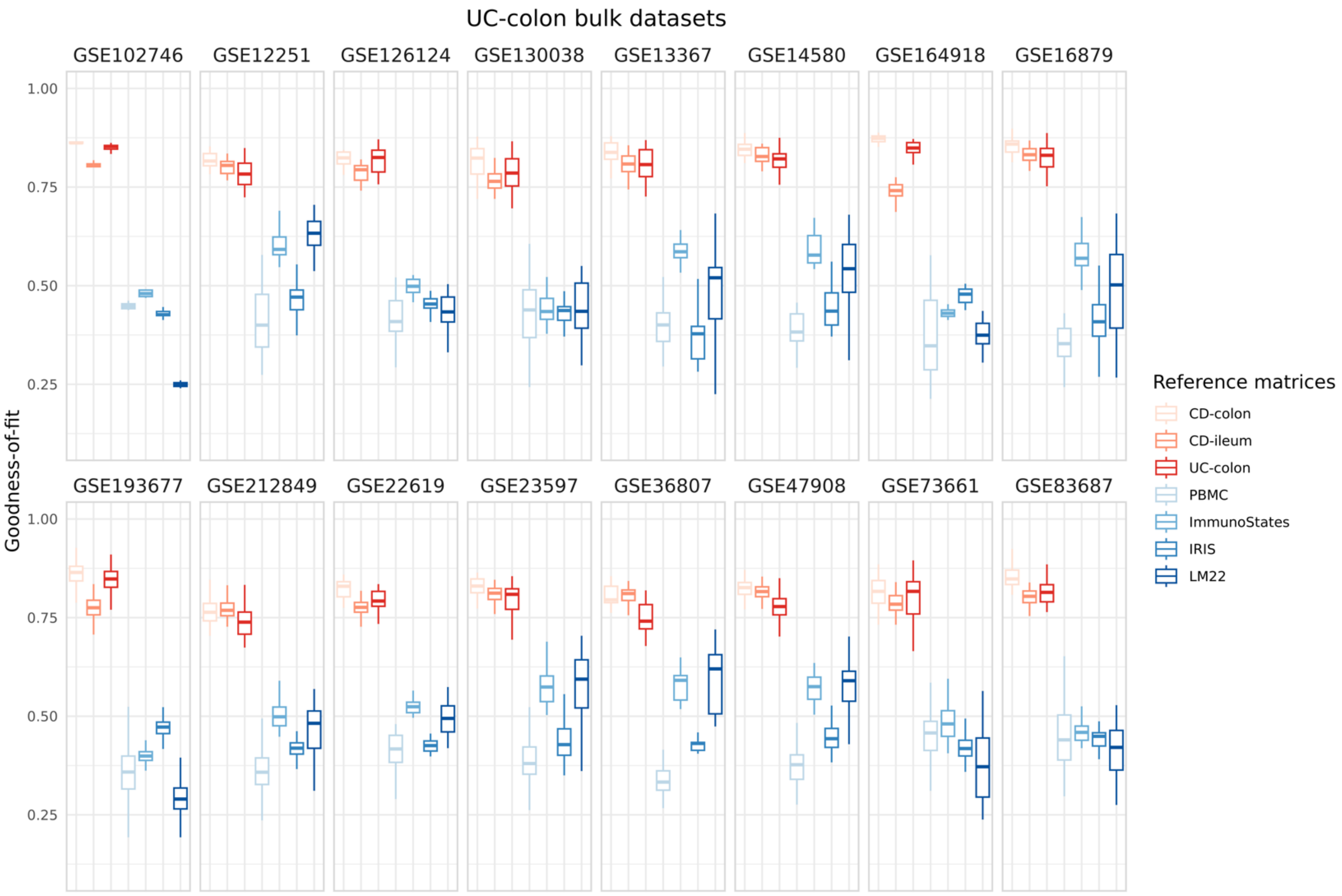


Fig. S3 Deconvolution performance on 16 public UC-colon bulk transcriptomic datasets.

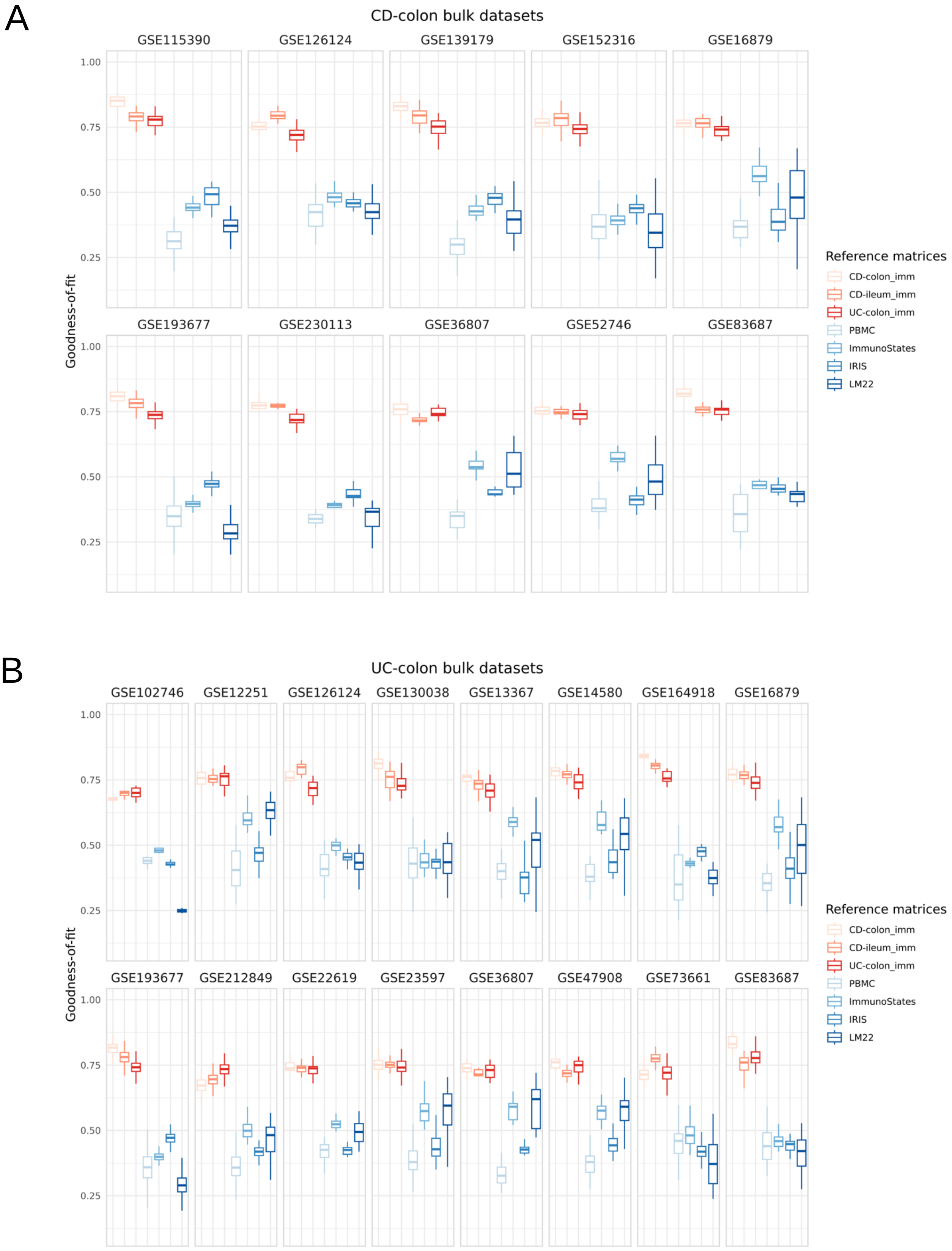


Fig. S4 Deconvolution performance on public (A) CD-colon (B) UC-colon bulk transcriptomic datasets using CRMs that are restricted to immune cell types.

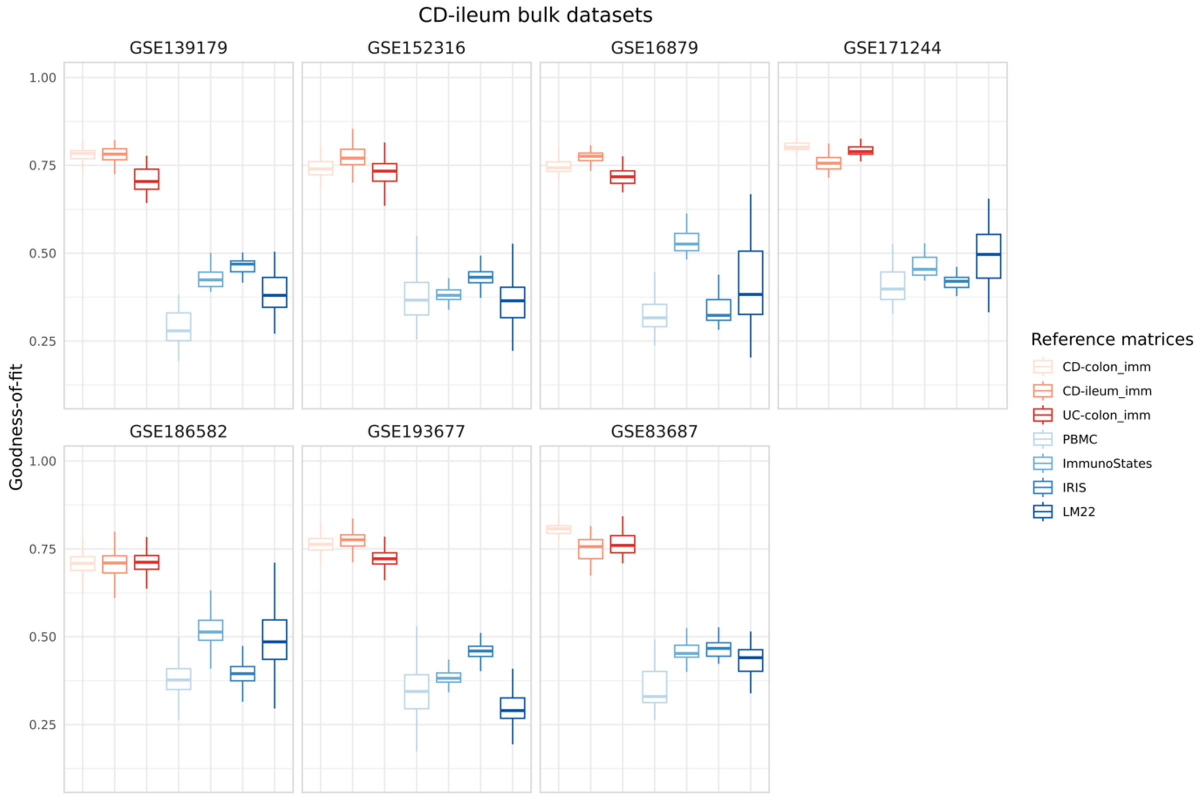


Fig. S5 Deconvolution performance on public CD-ileum bulk transcriptomic datasets using CRMs that are restricted to immune cell types.

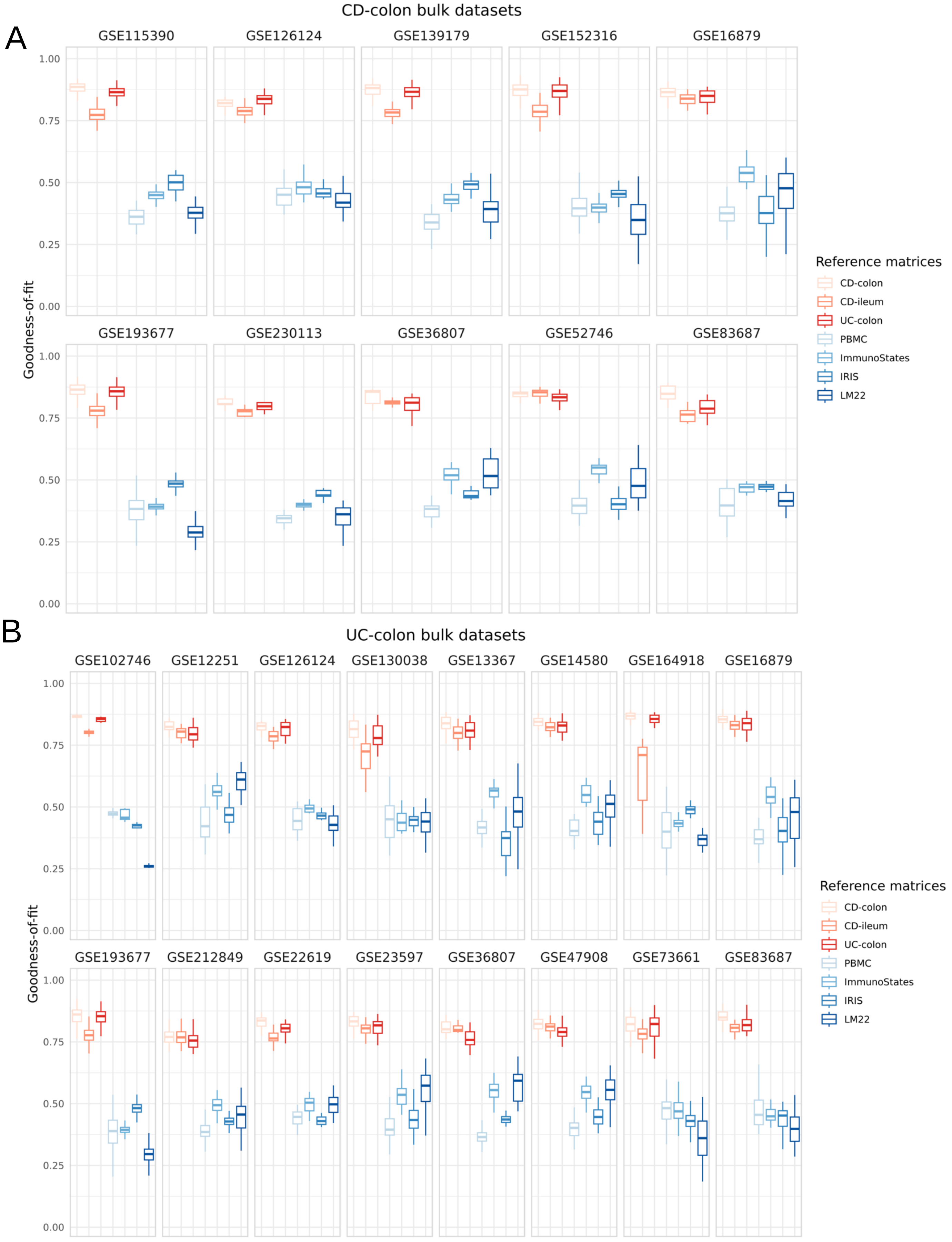


Fig. S6 Deconvolution performance on public (A) CD-colon (B) UC-colon bulk transcriptomic datasets using robust fitting of linear models (RLM).

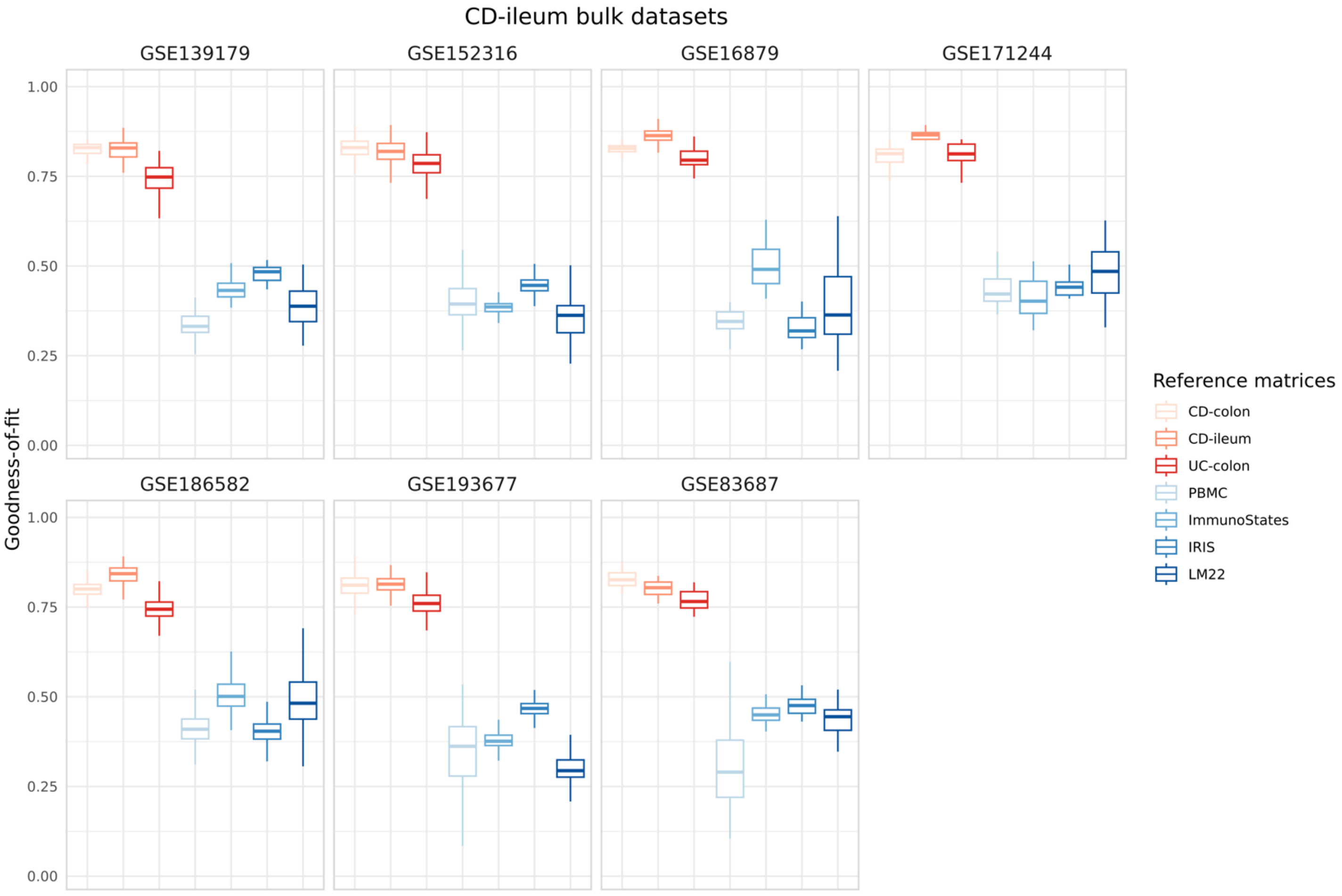


Fig. S7 Deconvolution performance on public CD-ileum bulk transcriptomic datasets using robust fitting of linear models (RLM).

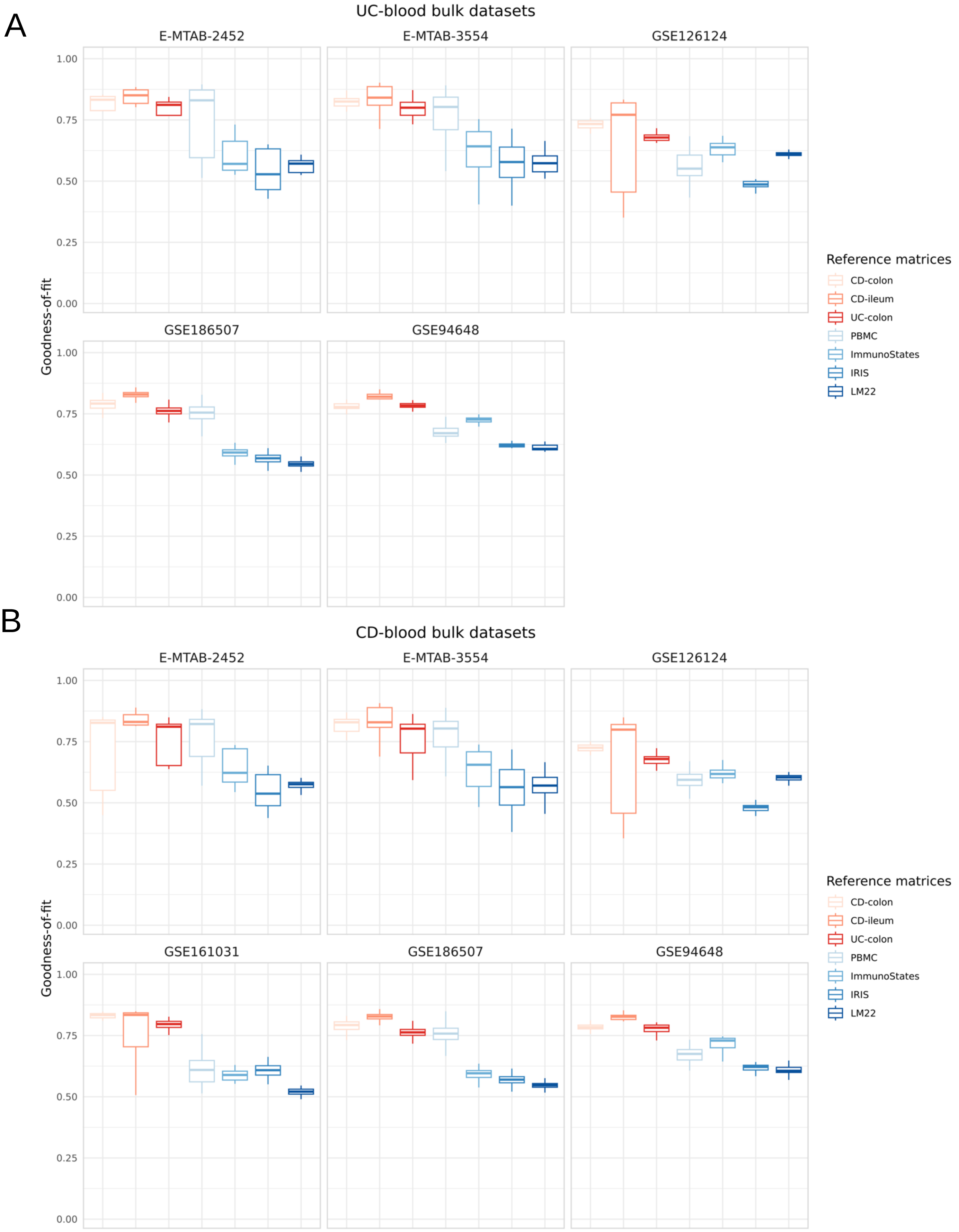


Fig. S8 Deconvolution performance on public (A) CD-blood (B) UC-blood bulk transcriptomic datasets using robust fitting of linear models (RLM).

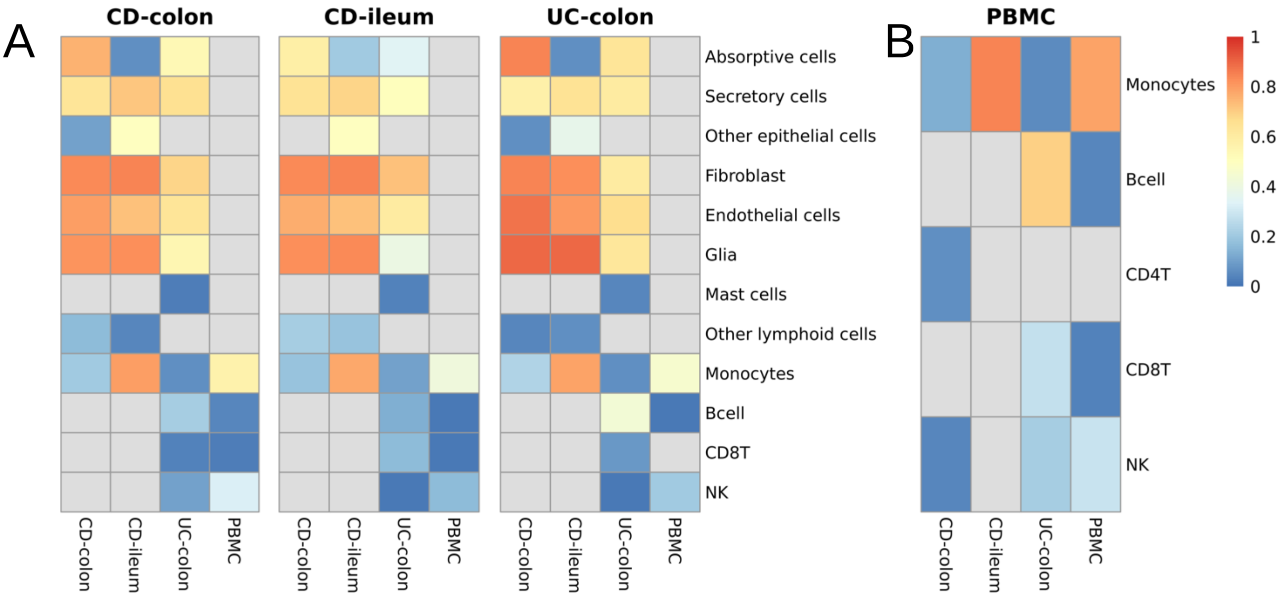


Fig. S9 The correlation of cell lineage proportions on simulated (A) tissue and (B) blood pseudobulk datasets using MuSiC for deconvolution.
